# Spatially-resolved integration of microglia morphological diversity and gene expression using Visium with protein co-detection

**DOI:** 10.64898/2026.06.05.730425

**Authors:** Jennifer Kim, Stephanie C. Hicks, Kristen R. Maynard, Paul Pavlidis, Annie V. Ciernia

**Author notes:** **Co-correspondence:** Annie V. Ciernia; Paul Pavlidis.

## Abstract

Microglia exhibit a dynamic range of morphologies that actively shift in response to local environmental cues. There has been a concerted effort as a field to move away from dualistic characterization of all microglia as ‘resting’ or ‘activated’, which is often described in terms of their morphological differences alone, towards more integrated definitions of microglial ‘states’ based on multiple data types. Recent advances in spatially-resolved transcriptomics offer new ways to understand microglial gene expression in situ. Here, we related microglia morphology to gene expression within a published, spatially-resolved proteogenomics (Visium-SPG) dataset of the human dorsolateral prefrontal cortex as proof of principle to combine the contributions of microglia morphology to regional transcriptomics differences in the brain. The published dataset uniquely combined spatial transcriptomics with immunofluorescent staining for four major brain cell types, including microglia, which enabled us to link microglial morphologies to gene expression in a spatially-resolved manner. Using a computational toolset, MicrogliaMorphology and MicrogliaMorphologyR, we classified individual microglia into morphological subtypes, assigned microglia to spots, and integrated these data with transcriptomic profiles for each Visium-SPG spot. We applied non-negative matrix factorization to account for contributions from multiple cell types, morphological features, and regional context in downstream differential expression modeling. Our approach offers a methodological framework for investigating the relationship between microglial morphology and gene expression in larger cohorts and within disease contexts where changes in microglia morphology are anticipated.

**Author summary:** Recent advances in spatially-resolved transcriptomics and the development of methods to deconvolve cell type signals has offered new ways to understand microglial gene expression in various tissue contexts. In this work, we related microglia morphologies to gene expression within a published spatial transcriptomics dataset of the human dorsolateral prefrontal cortex to assess whether morphologically different microglia are transcriptomically different in the healthy human brain. This dataset uniquely combines spatial transcriptomics with immunofluorescent staining for four major brain cell types, including microglia, enabling the linkage of microglial morphologies to gene expression in a spatially-resolved manner. Using a computational toolset for microglia morphology analysis, we classified individual microglia into morphological subtypes and integrated these data with spatial transcriptomic profiles while accounting for contributions from multiple cell types, morphological features, and regional context. This allowed for the identification of microglia morphology-related genes that are differentially expressed between microglial subtypes and relevant to human brain disorders. Findings together provide proof of principle for exploring the relationship between microglial morphology and gene expression in larger cohorts and diverse experimental contexts.

## Introduction

Clinical and postmortem studies(12–20) point to a neuroinflammatory component in the pathogenesis of brain disorders driven by enhanced activity of microglia, the brain’s resident macrophages. Single-cell or single-nucleus RNA sequencing (sc/snRNAseq) studies have shed new light on the various microglial states that exist across normal and diseased conditions and have yielded novel insights into the molecular mechanisms that shape a variety of context-dependent microglial responses.(21–25) However, these technologies require isolating cells from tissues, resulting in the loss of morphological information and spatial context. Microglia are highly dynamic cells that continuously sense and respond to cues in their microenvironments and rapidly change their morphologies, displaying various forms, which include ameboid, hypertrophic, rod-like or elongated, and ramified shapes.(26–30) These morphological changes are believed to play essential roles in their functions in health and disease, yet the relationship between microglial morphology and transcriptomic states remains elusive. While a diversity of microglial transcriptomic states are starting to be temporally, spatially, and context-dependently characterized, a gap exists in linking these states to other layers of information such as morphology to further probe their relevance to microglial functions.

Human snRNAseq studies suggest that different subsets of microglia contribute to specific disease mechanisms and that microglia can shift from homeostatic states to a variety of different inflammatory and disease-associated states in the progression of neurodegenerative disorders including Parkinson’s disease, Multiple Sclerosis, and Alzheimer’s disease (AD).(25,31–33) Changes in microglial morphology have been well documented in both mouse models of AD and postmortem human AD brains.(34–36) Microglia in the proximity of amyloid-beta plaques (Aβ) and tau-positive neurofibrillary tangles (tau), two pathological hallmarks of AD in the brain, exhibit reductions in arborization and branching complexity compared to microglia distant from the pathology. In the aging dorsolateral prefrontal cortex (DLPFC), microglia gene expression programs implicated in the accumulation of tau pathology are positively correlated with higher proportions of less branched microglia and higher tau accumulation.(37) While microglia spatially exhibit different morphological forms in relation to disease pathology(34–37), it is still unclear how coordinated changes in gene expression are related to these morphological changes in the healthy brain and in the transition to diseased states.

Recent technological advances in spatially-resolved transcriptomics (SRT) which couple gene expression and imaging of proteins offer promising avenues to probe these relationships using human brain tissue. In SRT technologies such as the Visium platform from 10X Genomics, tissue sections are mounted onto glass slides containing spatially arranged spots of a fixed diameter. Each spot contains barcoded capture probes that bind and collect mRNA from the tissue, enabling transcriptomic profiling at the spot level and preservation of spatial information. Chen et al.(38) was among the first to apply SRT technologies to study the spatiotemporal dynamics of microglia in the context of brain disorders, identifying gene signatures of microglia crosstalk with other brain cell types in proximity to AD pathological hallmarks. Several additional studies have profiled the spatial and anatomical distribution of microglial gene expression in relation to disease-relevant transcriptomics signatures.(39–42) However, few studies have successfully combined spatially-resolved transcriptomics(8,43) or proteomics(44) using immunohistochemical techniques to allow for high-throughput sequencing data to be linked to histologically defined microglial areas. While some efforts have been made to link microglial morphology to gene expression within the same samples, the focus was on individual morphological features, which ultimately provide incomplete representations of distinct morphological classes which vary along multiple axes including ramification, territory span, circularity, and branch length(29). To address these limitations, we developed new tools for relating SRT data to distinct microglia classes defined by multiple morphology measures.

As proof-of-principle for our approach, we leveraged a recently published spatially-resolved proteogenomics (Visium-SPG) dataset of the human DLPFC(8) to integrate microglia morphology with spot-level transcriptomics data. In the Visium-SPG dataset, the same brain sections used for spatial transcriptomics were first immunofluorescently stained for brain cell types, including microglia, and imaged prior to library preparation. The coupling of immunofluorescent (IF) signals with gene expression uniquely allowed us to apply our tools to integrate individual microglia morphologies with their respective transcriptomic profiles in a spatially-resolved and unbiased manner without the need for prior identification of morphology-related genes. We first classified individual microglia into *k*=4 morphological subtypes using a recently developed computational toolset, MicrogliaMorphology and MicrogliaMorphologyR(29), and integrated the resulting morphological data with the transcriptomics data for each barcoded Visium spot using custom ImageJ macros. Integration of these data types allowed us to identify and explore differentially expressed genes (DEGs) between morphological forms. In our DE modeling, we employed a spatially-aware approach using non-negative matrix factorization (NMF) to consider contributions of multiple brain cell types, unique ImageJ-derived spot-level features such as the proportion of spot area taken up by the microglia, and the regional makeup of the spots in our analysis. To facilitate future expansion of our work to other experimental contexts, we have provided all analysis code and ImageJ macros as free and open source resources. Our study provides proof of principle for similar work in the context of human brain disorders and can be extended to larger cohorts to better understand the relationship between microglial form and function.

## Materials and Methods

### Microglia morphology analysis of Visium-SPG images

To identify microglia morphological subtypes, TMEM119 channel .tiff images from Huuki-Myers et al., 2024(8) were used as input for analysis in MicrogliaMorphology and MicrogliaMorphologyR, as detailed in Kim et al., 2024a(29). Autofluorescence signal from respective lipofuscin channel .tiff images was first subtracted from the TMEM119 .tiff images, which resulted in images with lower background and true TMEM119 signal separated from background lipofuscin autofluorescence. Dataset-specific parameters used for thresholding included: (i) application of the Unsharp Mask filter using a radius of 20 and mask weight of 0.70, (ii) the Gaussian Blur filter using a sigma value of 1, and (iii) auto thresholding using the Default method for pixel values above 0. The Despeckle and Remove Outliers steps in the original protocol(29) were not included in the workflow for this dataset due to high signal to noise ratios in these images. An area filter of 1220-5728 um^2 was used for Step 3 of MicrogliaMorphology. After performing principal components analysis (PCA) on the 27 morphology features obtained from MicrogliaMorphology, the first 3 principal components (PCs) were used as input for fuzzy K-means clustering to define *k*=4 distinct classes of microglia morphology (ameboid, hypertrophic, elongated, ramified) with *k*-means cluster membership scores. The featurecorrelations(), pcfeaturecorrelations(), and clusterfeatures() functions from the MicrogliaMorphologyR R package(29) were applied to the data to explore the relationships between the morphology features in the dataset, to describe how the first 3 PCs explain the variability in the morphology features, and to assign cluster labels for the morphological forms. To generate Fig. 1G, the clusterfeatures() function was used for comparative analysis of clusters across the Visium-SPG morphology data and published mouse data (MicrogliaMorphologyR::data_2xLPS_mouse_fuzzykmeans). The 27 morphology features were scaled within datasets prior to hierarchical clustering to generate Fig. 1G.

**Figure 1.**
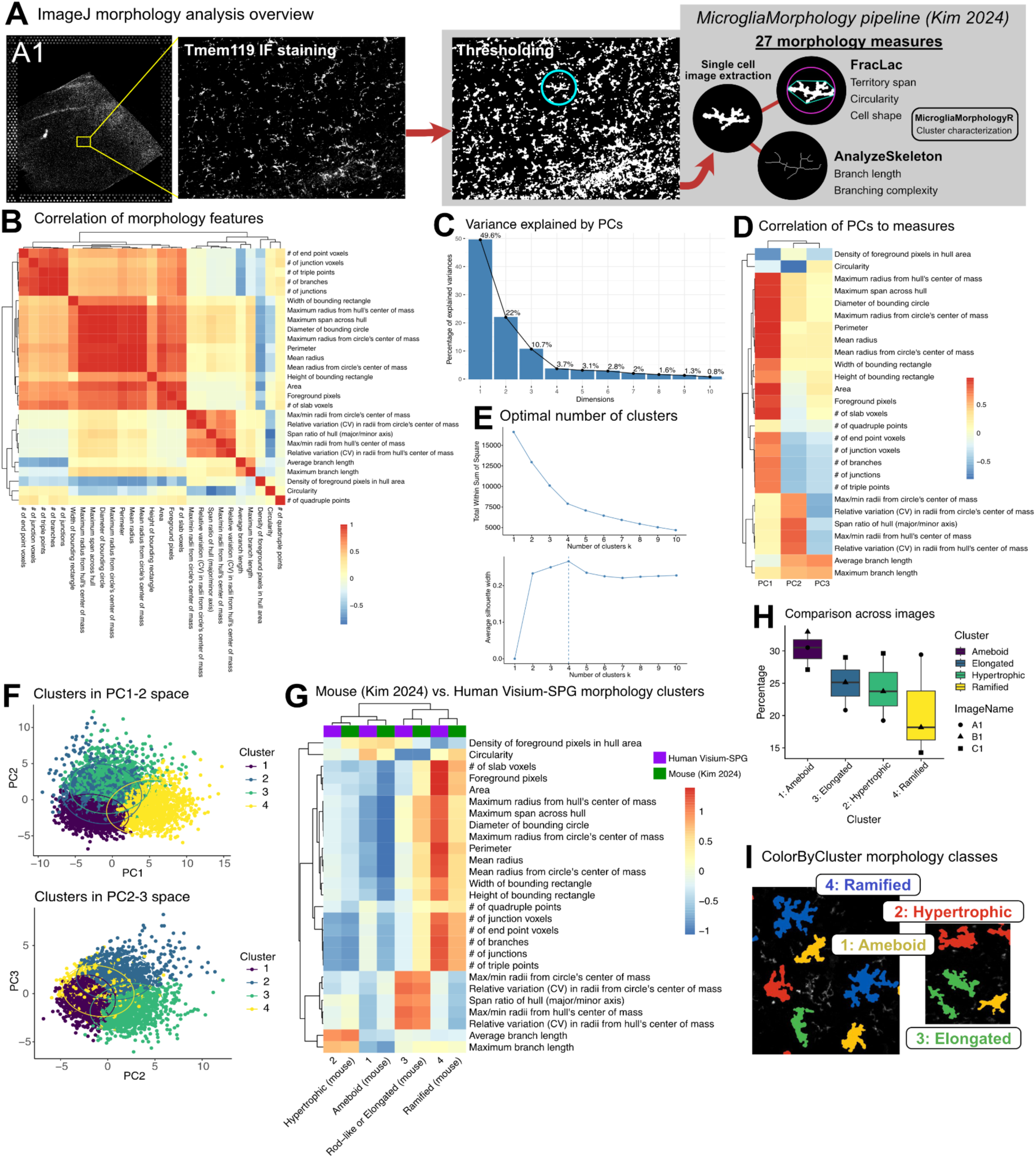
Identifying ameboid, hypertrophic, ramified, and elongated microglia morphologies using MicrogliaMorphology and MicrogliaMorphologyR. (**A**) Overview of morphology analysis (n=3 images from Huuki Myers et al., 2024; Image A1 = Br2720_Ant_IF, 2668 cells; B1 = Br6432_Ant_IF, 1293 cells; C1 = Br6522_Ant_IF, 1583 cells). (**B**) Spearman’s correlation of 27 morphology features output by MicrogliaMorphology. (**C**) Variance explained by the first 10 PCs after dimensionality reduction on 27 morphology features in A. (**D**) Spearman’s correlation of morphology features to the first 3 principal components. (**E**) Optimal k-means clustering parameters determined using total within sum of squares and average silhouette width techniques. (**F**) Data points displayed in PCs 1-2 and 2-3 space, colored by k-means cluster. (**G**) Comparative analysis of morphology clusters between published mouse data (Kim et al., 2024) with Visium-SPG human clusters. Mean values for all 27 morphology features, scaled within datasets prior to hierarchical clustering. (**H**) Percentage of each morphology cluster present in each Visium-SPG image analyzed. (**I**) Morphology classes visualized using ColorByCluster feature within MicrogliaMorphology pipeline.

### ImageJ integration of Visium-SPG dataset with MicrogliaMorphology output

#### Generation of final microglia ROIs and Visium spot ROIs

The CellIDs of the final microglia that were included in MicrogliaMorphologyR analysis were saved as one column in .csv files for each image. A custom macro was used (‘SavingJustCellROIs.ijm’) to generate collections of final cell ROIs for each image using these .csv files and thresholded versions of the images as input. A custom macro was used (‘VisiumSpotOverlay.ijm’) to generate image-specific Visium spot ROIs. Visium spot coordinates (x, y) for each image (A1, B1, C1) from the published dataset were loaded as tables into ImageJ and the makeOval() function was used to create circles of image-specific diameters available with the dataset (A1=55.347, B1=55.3147, C1=55.3153) for each barcoded spot. A collection of image-specific Visium spot ROIs were saved for downstream analysis.

#### Analysis of fluorescent channel signals at Visium spot level

Autofluorescence signal from lipofuscin channel .tiff images was first subtracted from the channel-specific .tiff images. Image-specific Visium spot ROIs were loaded in ImageJ and overlaid on top of each fluorescent channel .tiff image for downstream measurements of TMEM119, GFAP, NEUN, and OLIG mean signal intensity and area within the spots using the ROI manager. (Fig. S1A-C) For each of the brain cell types in the dataset, Pearson’s correlations were run across the channel-specific ImageJ-derived measures (mean signal intensity or signal area) and the cell type proportions and numbers estimated by deconvolution methods provided in the Visium-SPG dataset (Cell2Location, Tangram, CART). (Fig, S1D)

#### Area overlap analysis to generate ‘proportion_spotarea’ and ‘proportion_cellarea’ measures

For each image analyzed, the thresholded image, final cell ROIs, and Visium spot ROIs were loaded into ImageJ in this order before running a custom macro (‘AreaOverlap.ijm’) to calculate the overlapping areas between the microglia cell ROIs and Visium spot ROIs. The resulting output included measurements from only those cells that overlapped with Visium spots. The outputs for each image were saved as .csv files and loaded into R, where the percentage of the microglia cell area within a Visium spot (proportion_cellarea = area overlap / cell area) and the percentage of spot area occupied by a microglia cell (proportion_spotarea = area overlap / spot area) were calculated.

### Marker gene expression analysis: fluorescent channel signals (Fig. S2A)

Marker gene lists were downloaded from PanglaoDB(7) and filtered for microglia, neuron, astrocyte, and oligodendrocyte cell type-markers. The Visium gene expression dataset was filtered to include only genes from the marker gene lists. Within each image, for each cell type-specific channel (TMEM119: microglia, NEUN: neurons, GFAP: astrocytes, OLIG: oligodendrocytes), correlations between ImageJ measures (mean signal intensity or total signal area) and expression of cell type-specific marker genes were calculated for each Visium spot in the image. Each marker gene’s correlation to the respective measures was represented as a single point in violin plots generated using the ggplot()(45) function in R. (Fig. S2A)

### Pseudobulk sample generation

Prior to pseudobulking, the following filtering criteria were applied in this order to isolate Visium spots that included the most representative population of distinct microglia morphology classes as possible (Fig. 2A):

1. Filtered to include microglia cells with one Visium spot assignment (i.e., microglia cells do not span more than one Visium spot)
2. Filtered to include the Visium spots that have only one microglia morphology class present (i.e., spot can have 2+ microglia as long as they belong to the same morphology class)
3. Filtered to include Visium spots that have 10% or more of the microglia cell in the spot (proportion_cellarea ≥ 0.10)
4. Filtered to include microglia with a 50% or more fuzzy *k*-means membership score for a given cluster to examine the most representative cells for each class.

**Figure 2.**
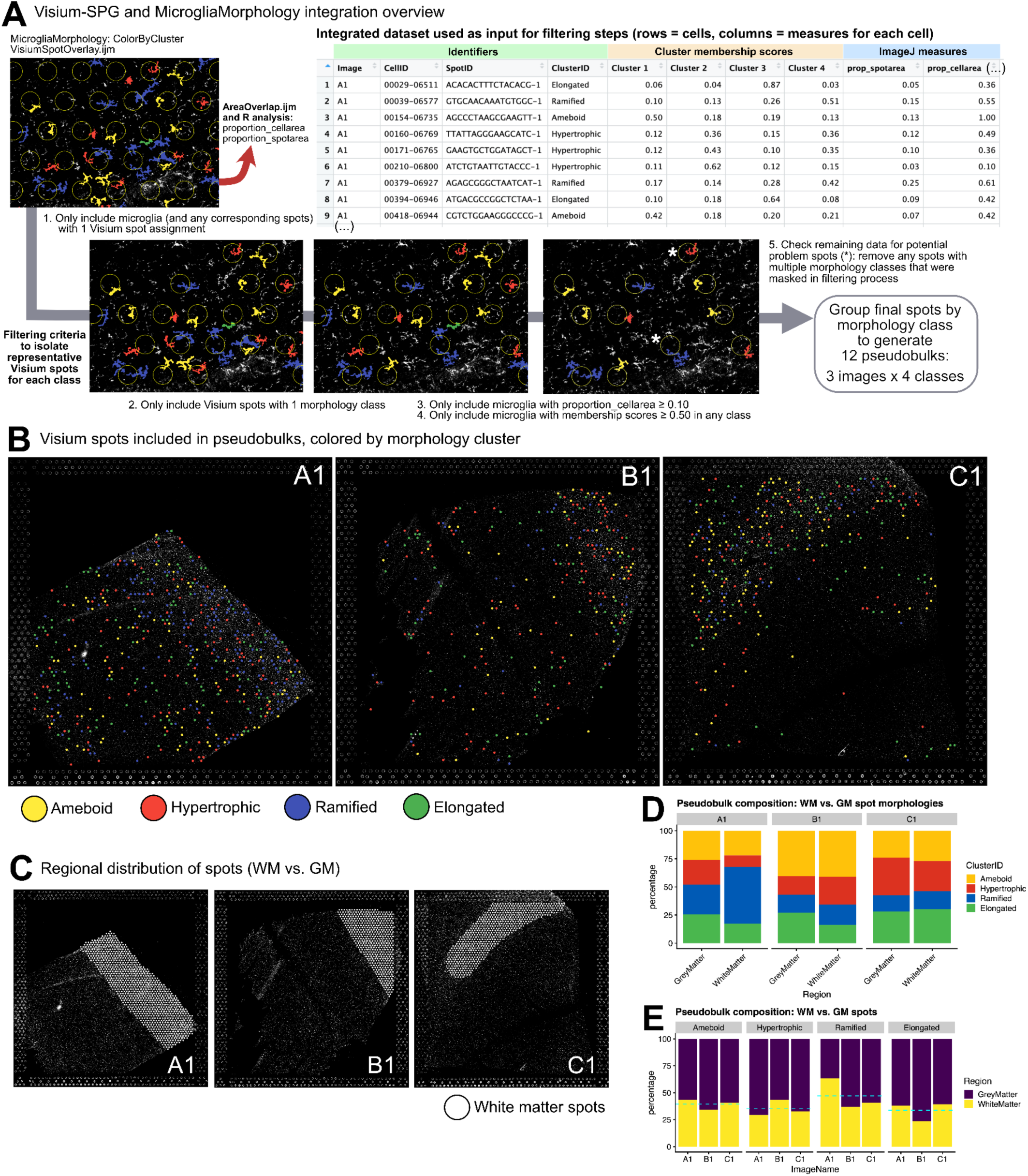
Integration of MicrogliaMorphology output with Visium gene expression data. (**A**) Overview of how Visium-SPG and MicrogliaMorphology data were integrated. (**B**) Visualization of final Visium spots included in generation of pseudobulks. (**C**) Visualization of white matter spots in Visium-SPG dataset. (**D**) Composition per image of spot morphologies in grey vs. white matter. (**E**) Composition per pseudobulk of grey vs. white matter spots.

As a final step to stay faithful to our filtering criteria, the remaining Visium spots were examined to ensure that they truly had one microglia morphology class present, as this could have been masked when filtering in Steps 1-4 (See Fig. 2A, S3A-B for example visualization of potential problem spots that were further examined). A total of 8 Visium spots were removed at this step (Fig. S3B).

We created 12 pseudobulked samples (3 images x 4 morphology clusters/image) from the post-filtering, which were used as input samples for differential expression (DE) analysis. Counts were aggregated (summed) across spots for each gene to generate pseudobulk profiles. Each pseudobulked sample contained the following numbers of spots:

The ImageJ-derived variables for each pseudobulked sample were aggregated for the spots included in each sample: a) pseudobulked proportion_spotarea = aggregated spot area taken up by cell / (number of visium spots x π x image-specific spot radius ^2) and b) pseudobulked proportion_cellarea = aggregated cell area that made it into spot / aggregated total cell areas. The proportion of white matter spots included in the pseudobulked sample (proportion_WM), was calculated using the spot layer annotations (‘manual_layer_labels’ metadata column) included in the Visium-SPG dataset(8). CART cell type proportions were also calculated at the pseudobulk level by aggregating counts for each cell type across the spots included for each sample. Spearman’s correlations were calculated across variables present in the pseudobulked dataset including CART cell type proportions (microglia, neurons, astrocytes, oligodendrocytes, other cell types), ImageJ-derived measures (proportion_spotarea, proportion_cellarea), and a region composition measure (proportion_WM).

### Marker gene expression analysis: pseudobulked spots (Fig. S2B-C)

The marker gene list downloaded from PanglaoDB(7) was filtered for liver cells to use as a negative control gene list. The final Visium spots that were included in the 12 pseudobulked samples detailed in Table 1 were either grouped as a single sample (n=1104 spots) or split into 2 region-specific samples for white matter (n=456) and grey matter (n=648) spots using the spot layer annotations (‘manual_layer_labels’ metadata column) included in the Visium-SPG dataset(8). Counts (‘spaceranger count’ output included with Visium-SPG dataset) across spots included in the pseudobulked samples were aggregated for each gene, and only genes with at least 1 count were included in analysis. Genes were then stratified into low, medium, and high expressing categories by dividing their log(count) values into tertiles using the ntile() function in R (Fig. S2B-C). For each sample (all spots, white matter spots, grey matter spots), gene overlap enrichments between low, medium, and high expressing genes and PanglaoDB marker gene lists (brain cell types: microglia, neurons, astrocytes, oligodendrocytes; liver cell types as a negative control) were calculated using the newGOM() function in the GeneOverlap() R package(46). For each of the 3 grouped samples (all spots, WM spots, GM spots), the genome.size (or universe) argument was set as the final numbers of genes included that were specific to each sample after filtering genes with low read counts. Fisher’s exact tests were applied and overlap enrichment between lists was represented as Odds Ratios with BH-adjusted *p*-values (Fig. S2B-C).

**Table 1.** Total Visium spot numbers for each pseudobulked sample.

| Pseudobulk ID | ImageName | Morphology cluster | Number of Visium spots |
| --- | --- | --- | --- |
| A1_1 | A1 | Ameboid | 122 |
| A1_2 | A1 | Hypertrophic | 82 |
| A1_3 | A1 | Elongated | 110 |
| A1_4 | A1 | Ramified | 191 |
| B1_1 | B1 | Ameboid | 111 |
| B1_2 | B1 | Hypertrophic | 53 |
| B1_3 | B1 | Elongated | 64 |
| B1_4 | B1 | Ramified | 46 |
| C1_1 | C1 | Ameboid | 81 |
| C1_2 | C1 | Hypertrophic | 101 |
| C1_3 | C1 | Elongated | 94 |
| C1_4 | C1 | Ramified | 49 |

### Non-negative matrix factorization

Input variables (CART cell type proportions: microglia, neurons, astrocytes, oligodendrocytes; ImageJ-derived features: proportion_cellarea, proportion_spotarea; and region composition feature: proportion_WM) were min-max transformed to keep all variables within the same scale (0-1). The reconstruction errors and percentage variance explained were calculated for a range of 1-10 components (Fig. 3C) to determine the optimal number of components within this range to include in downstream analysis. Non-negative matrix factorization (NMF) dimensionality reduction was run for 4 components using the “brunet” method by applying the nmf() function within the NMF R package(47). The NMF component loadings for each pseudobulked sample were extracted from the dataset using the basis() function and input as variables for the final statistical models used in differential expression analysis.

**Figure 3.**
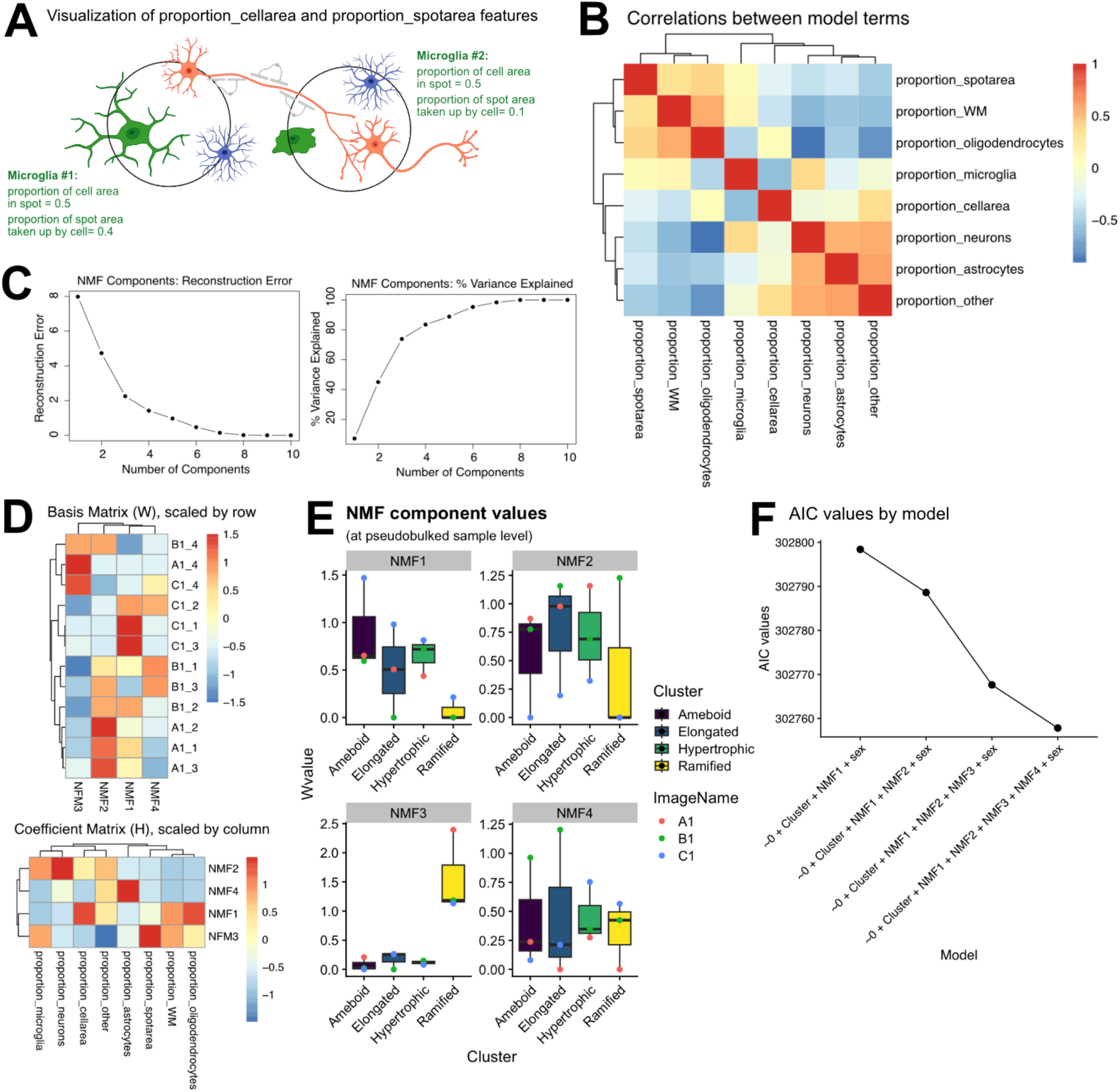
Final differential expression model was fit to components derived from non-negative matrix factorization. (**A**) Visual example of proportion_cellarea and proportion_spotarea ImageJ-derived features. Example microglia are in green, neurons in red, astrocytes in blue, and oligodendrocytes in grey. Made with Biorender. (**B**) Spearman’s correlation of ImageJ-derived features and CART cell type proportions. (**C**) Optimal number of NMF components to include determined using reconstruction error and percentage cumulative variance explained. (**D**) NMF basis matrix scaled across components for each pseudobulked sample and NMF coefficient matrix scaled across components for each input feature. (**E**) NMF component values between morphological classes across images. (**F**) AIC values calculated for the eBayes fit of each model specific to DE contrasts (pairwise, 1 vs. all others; 10 total, see Fig. 3B-C).

### Differential expression analysis

Low expressing genes were removed by filtering to include genes with at least 1 count per million (CPM) in at least 1 sample (Fig. S4A). The remaining genes (n = 15,137) were normalized using the Trimmed Mean of M-values (TMM) method to correct for library composition using the calcNormFactors() function in the edgeR R Package(48) (Fig. S4B). Dimensionality reduction by Principal Components Analysis (PCA) was applied to evaluate the contribution of ImageName and Morphology Cluster to the overall gene expression dataset (Figs. S4C-D). To determine how many NMF components to include in the final differential expression (DE) model, Akaike Information Criterion (AIC) values for each model (varying the number of NMF components to include) were calculated to test the goodness of each model’s fit to the data (Fig. 3F). AIC values were calculated for the eBayes fit of each model specific to the DE contrasts (10 total), which included 1 vs. all contrasts (e.g., hypertrophic vs. all other morphology clusters) and pairwise contrasts (e.g., ramified vs. ameboid). Filtered CPM values were input into the voom() function within the limma R package(49) using the final DE model: ∼0 + Cluster + NMF1 + NMF2 + NMF3 + NMF4 + sex, specifying ImageName as a random effect (block=ImageName). Individual contrast comparisons were then called using contrasts.fit() followed by eBayes(). Differentially expressed genes (DEGs) were identified for each comparison of interest using limma’s topTreat() function with a BH *p*-value correction (significance at q < 0.05 and fold change > ± 2).

### Gene enrichment analysis

Gene set enrichment was performed on DEGs from the Ameboid vs. All contrast using the gsr() function from the R package ErmineJ(1–3). Fold changes were used as gene scores (Fig. 4B). Gene list enrichment was performed against published microglia lists using the 2025 database of the MGEnrichment app(4), with all genes interrogated in the experiment as background. Significant enrichments against MGEnrichment gene lists were tested by a one-tail Fischer’s exact test and BH-corrected to reach significance at an adjusted p-value < 0.05. Target % and Background % values for significant gene list enrichments were plotted using ggplot2 in R (Fig. 4C).

**Figure 4.**
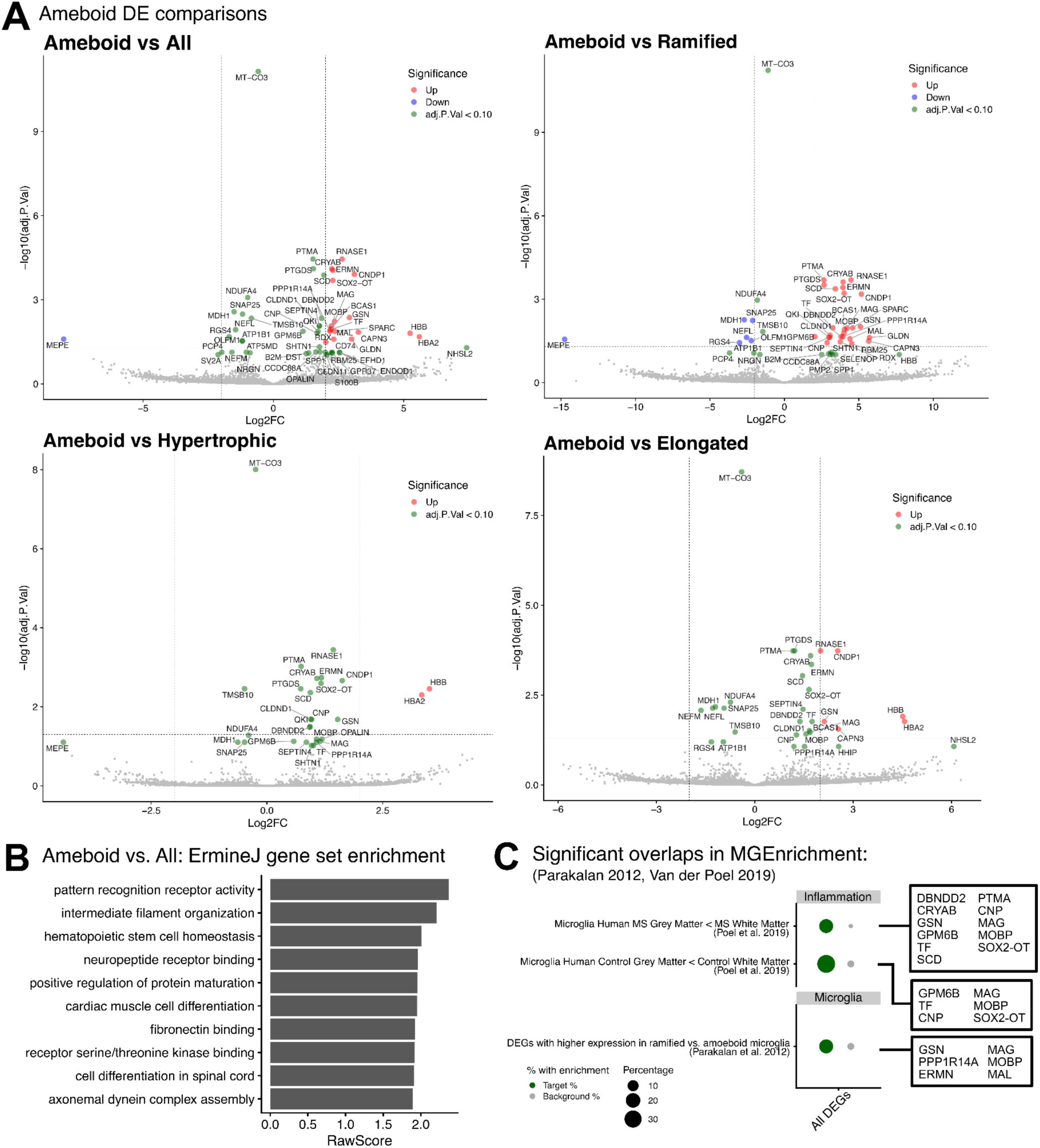
Example differential expression analysis. (**A**) Volcano plots of Ameboid DE comparisons (ameboid vs. all other classes, pairwise) (* q<0.05, log2FC> +/-2). (**B**) ErmineJ(1–3) gene set enrichment for Ameboid vs. All DE list using fold changes as gene scores. (**C**) Gene list overlap enrichment against publicly available microglia lists in MGEnrichment(4) and all 34 significant DEGs (across all comparisons including ameboid contrasts). Green circles denote a significant enrichment in the percent of DEGs for a given gene list compared to that of the background of all genes detected in the RNA-seq experiment (grey circles). Only significant enrichments are shown (BH-corrected p<0.05). Of the significant enrichment results, only displaying those from Van der Poel et al., 2019(5) and Parakalan et al., 2012(6). All other significant enrichments can be found in Fig. S6.

### Exploration of spatial and single-nucleus expression of under-threshold DEGs (SPP1, CD74)

The vis_gene() function within the spatialLIBD R package(50) was used to plot the spatial, spot-level expression of *SPP1* and *CD74* for each of the images analyzed in our dataset (Image A1 = Br2720_Ant_IF, B1 = Br6432_Ant_IF, C1 = Br6522_Ant_IF) (Fig. S5A). Paired snRNAseq data from Huuki-Myers et al., 2024(8), which included the 4 individuals represented in the Visium-SPG dataset, was downloaded using the spatialLIBD R package(8). The snRNAseq dataset was generated from adjacent DLPFC tissue blocks from 30 healthy adult donors within the same study. The ggcells() function within the scater R package(51) was used to create boxplots of cell type-specific gene expression for each of the genes using the annotations available in the ‘cellType_broad_hc’ column within the snRNAseq SingleCellExperiment(52) R object (Fig. S5C). Cross-species expression of the genes were plotted by brain cell type using publicly available gene expression datasets of purified brain cell type populations from mouse and human brain that were downloaded from BrainRNASeq.org(9,10) (Fig. S5D). Microglia cluster-specific expression of the genes within the publicly available Human Microglia Atlas(11) was plotted using the scCustomize(53) R package. (Fig. S5E)

## Results

### MicrogliaMorphology and MicrogliaMorphologyR can be used to identify ameboid, hypertrophic, ramified, and elongated microglia morphological forms in Visium-SPG data

To characterize and correlate microglia morphology to gene expression, we leveraged a publicly available spatially-resolved proteogenomics dataset (Visium-SPG, 10X Genomics), Huuki Myers et al., 2024(8), from the dorsolateral prefrontal cortex (DLPFC) of 3 neurotypical humans. This dataset included immunofluorescent images for TMEM119, NEUN, GFAP, and OLIG2 to label microglia, neurons, astrocytes, and oligodendrocytes, respectively, from the same DLPFC tissue sections that underwent spatial transcriptomics profiling. We used MicrogliaMorphology to extract 27 morphology features from 5544 individual microglia cells in the TMEM119 channel images (Fig. 1A). Within the feature space, we identified distinct sets of highly correlated morphology features describing branching complexity, territory span, cell shape, and branch length (Fig. 1B), as previously characterized(29,30). The first 3 principal components (PCs) after PCA described 82.3% of the variability in the dataset (Fig. 1C). Relative to the other PCs used for clustering, PC1 (49.6%) mostly described features related to territory span, PC2 (22.0%) to cell shape and circularity, and PC3 (10.7%) to branch length (Fig. 1D). PCs 1-3 were used as input for fuzzy K-means clustering. Using a soft clustering approach, fuzzy k-means scores allowed us to isolate the Visium spots containing the most representative cells of each morphology class for downstream analysis. The total within sum of squares and average silhouette metrics (Fig. 1E) were used to determine an optimal clustering parameter of *k*=4 microglia morphological forms (Fig. 1F). Broadly, clusters that had the highest relative territory span and branching complexity values were labeled as ‘ramified’, those that had the highest cell oblongness as ‘elongated’, those with the highest circularity and pixel density with the lowest territory span as ‘ameboid’, and those with the lowest branching complexity as ‘hypertrophic’. These labels corresponded to morphology clusters previously characterized in a published mouse dataset(29) (Fig. 1G). Consistent with previous findings in human brain studies, all four microglia morphologies were present in significant proportions(54). We observed similar proportions of each morphology cluster across images (group means: hypertrophic, 24.2%; ameboid, 30.2%; ramified, 20.6%; elongated, 25%) (Fig. 1H). Cluster labels were visually verified using the ColorByCluster feature within MicrogliaMorphology (Fig. 1I). Taken together, we identified 4 morphologically distinct clusters in the human DLPFC that were correlated with published mouse microglia morphologies.

### Cell type-specific correlations of immunofluorescent signals with gene expression and exploration of grey vs. white matter contributions to pseudobulked samples

To evaluate the consistency of the staining panel across images, we assessed the staining quality of each cell type marker. Visium spots (55-µm diameter) were overlaid onto each fluorescent channel image (GFAP, NEUN, OLIG2, TMEM119) so that ImageJ-derived features, including ‘mean signal intensity’ and ‘signal area,’ could be calculated for each channel across the entire Visium capture area as well as at the individual spot-level. Channel signal intensity within the entire Visium capture area was similar across images for each cell type marker (Fig. S1A). When assessing at the spot level, compared to the other images in the dataset, Image D1 (‘Br8667_Post_IF’ in the publicly available dataset) had smaller signal area values (Fig. S1B) and mean signal intensity values (Fig. S1C) for TMEM119. These metrics were both centered near 0, indicating an abundance of punctate rather than full, connected microglial staining. Image D1 was thus considered to have microglial staining inadequate for morphological assessment and removed from downstream analysis.

As Visium spots are 55um in diameter, they contain multiple cell types and in order to parse out gene expression signatures unique to morphological microglia, we needed to estimate the contributions of other cell types in the spots to consider for DE analyses. There were several estimates available in the Huuki-Myers 2024(8) dataset: *Classification and Regression Tree (CART)(8)* estimates which are based on the immunofluorescent imaging data, and *Tangram(55)* and *Cell2location(56)* which infer cell types based on gene expression. To determine which estimates of cell type composition to use in downstream modeling, we compared our ImageJ-derived features to estimates that were available in the Visium-SPG dataset. Across brain cell types, the ImageJ features correlated most highly with the CART(8) approach-derived measures of cell type proportions compared to estimates derived from other cell deconvolution methods in the Visium-SPG dataset (Fig. S1D). CART estimates were thus used as estimates of cell type-specific proportions in downstream differential expression analyses. To explore how microglia fare in cell type-specific gene expression signals compared to other brain cell types within the Visium-SPG dataset, we analyzed the expression patterns of marker genes listed in PanglaoDB(7), a publicly available database of integrated sc/snRNAseq data from published mouse and human studies. Compared to other brain cell types, neurons had the highest concordance between cell type-specific marker gene expression and cell type-specific channel signals across images within the individual Visium spots (Fig. S2A).

To further assess whether we could detect microglial marker gene expression across the final spots included in our pseudobulked DE samples (Table 1, Fig. 2A-B, S3) and to evaluate whether there was any region-specificity in the ability to detect microglial signals, we created a pseudobulked sample which aggregated all spots regardless of region as well as region-specific pseudobulks based on spot location in white matter (WM) vs. grey matter (GM). We then compared the total, WM, and GM pseudobulked samples. Genes with at least 1 count in the pseudobulks were stratified into low, medium, and high expressing categories by dividing their log(count) values into tertiles. We found that neuron, astrocyte, and oligodendrocyte marker genes overlapped with the high-expressing genes (Fig. S2B-C; BH-adjusted *p*-values < 0.05). Microglia marker genes overlapped with the medium-expressing genes when all spots were pseudobulked together and when separated into GM, but not WM spots (Fig. S2B-C). This suggests that the cell type makeup of the WM vs. GM environments could potentially affect the ability to detect microglial signals. To account for this potential impact on gene expression driven by region-specific composition, we calculated a new variable called “proportion of pseudobulked spots in white matter” (proportion_WM) to include in downstream DE analysis (Fig. 2C,E). There was variability in the region-specific proportions of each morphology class for each image (Fig. 2B-D), but there were no striking differences on average in GM vs. WM spots included in the final morphology-specific pseudobulks included for DE analysis (Fig. 2E).

### ImageJ-derived features were integrated with Visium-SPG data to model microglia morphology-relevant gene expression signatures

Integration of the morphology and spot-level data allowed us to calculate novel spot-level measures, including the proportion of each cell within its respective spot (proportion_cellarea) and the proportion of each spot that was taken up by microglia (proportion_spotarea). (Fig. 3A) We considered these ImageJ-derived features, as well as the cell type proportions available in the Visium-SPG dataset, in downstream DE modeling on 12 total pseudobulked samples for each morphology class from each sample (Fig. S4). Correlations across the final variables calculated for the pseudobulked samples – CART proportions (microglia, neurons, astrocytes, oligodendrocytes, other cell types), proportion_cellarea, proportion_spotarea, and proportion_WM – revealed new relationships between the variables. For example, proportions of neurons, astrocytes, and other cell types were highly correlated to each other, but anti-correlated with proportion_oligodendrocytes, proportion_spotarea, and proportion_WM. (Fig. 3B).

Because Visium spots contain multiple cell types, we wanted to preserve the relationships between these variables and control for them when making downstream comparisons between our morphologically-defined pseudobulks. We applied dimensionality reduction by non-negative matrix factorization (NMF) on all of the variables and used the resulting NMF component values as the terms in DE modeling. Modeling on the NMF components allowed us to consider how cell type proportions fluctuate with respect to each other, how much physical space different microglia morphologies occupy within the sampled space, how much of the microglial cell was found in the spots assessed as a proxy for how much microglial mRNA was available for transcriptomics library preparation, and the regional makeup (WM vs. GM) of the pseudobulked spots. In this way, we were able to analyze gene expression differences between pseudobulked samples representative of distinct microglial morphologies while also accounting for additional factors that could contribute to the differences in gene expression between the samples. Using the reconstruction error, we determined a parameter of 4 components to include for the NMF analysis, which together described 83.5% of the variability in the dataset. (Fig. 3C) Each NMF component differentially described the variability in the dataset explained by different features. Compared to the other components, NMF1 was most highly correlated to proportion_oligodendrocytes, NMF2 to proportion_neurons, NMF3 to proportion_spotarea and proportion_other, and NMF4 to proportion_astrocytes. (Fig. 3D) The pseudobulked samples were differentially weighted for the NMF components individually (Fig. 3D) and when they were grouped by morphology class (Fig. 3E). AIC values, which test goodness of fit while penalizing for model complexity, were minimized when considering all four NMF components in the model, indicating that the final model was best fit to the data: ∼0 + Cluster + NMF1 + NMF2 + NMF3 + NMF4 + sex. (Fig. 3F)

### Integrative approach can be used to relate microglia morphology to transcriptomics differences in Visium-SPG data

34 total DEGs were identified across all comparisons, which included 1 vs. all contrasts (e.g., hypertrophic vs. all other morphology clusters) and pairwise contrasts (e.g., ramified vs. ameboid). Differential gene expression was mostly driven by the ameboid vs. all others and ameboid vs. ramified forms across comparisons (Fig. 4A; Supplementary Table 1; * q<0.05, log2FC> +/-2). Focusing on the ameboid vs. all comparison as an example, DEGs were enriched for terms relevant to immune function such as pattern recognition receptor activity, and cell motility and morphology such as intermediate filament organization and axonemal dynein complex assembly. (Fig. 4B; Supplementary Table 1) 11 of the total 34 DEGs across all comparisons overlapped with genes previously identified as upregulated in WM vs. GM-residing human microglia(5) and higher in expression in ramified vs. ameboid mouse microglia(6). (Fig. 4C; Supplementary Table 1) From our DE analysis, we identified two morphology-relevant targets previously recognized in the literature, *SPP1* and *CD74,* which have previously been identified as important markers of disease-associated microglial (DAM) states(25,25,57–59). Although both of these genes were under significance threshold (for ameboid vs. all others contrast, SPP1: log2FC=1.79, q=.07; CD74: log2FC=2.25, q=.07), they were 1) co-localized with morphologically segmented microglia (Fig. S5A-B), 2) highly expressed in microglia in the paired snRNAseq dataset(8) (Fig. S5C), which included the 3 Visium-SPG subjects, and 3) had microglia-specific expression in independent human(10) and mouse(9) single-cell atlases (Fig. S5D-E). Both *SPP1* and *CD74* were enriched in disease-associated microglia (DAM) clusters within a published Human Microglia Atlas(11). (Fig. S5E)

## Discussion

The recent advent of single-cell and spatial sequencing technologies has shifted the field towards a deeper understanding of microglial ‘state’ defined by various parameters such as gene expression, morphology, epigenetic regulation, biological context, and spatial distribution.(24,60) There now exists a plethora of methods that can be used together for multi-dimensional profiling of individual microglia, but it is unclear how these different approaches to defining state align with each other to produce a unified picture of microglia. While a range of different microglial morphologies have been shown to correspond with specific functions and disease states, transcriptomic distinctions across these morphologies have not yet been established. Previous work in rodent(6) and cell culture(61) models has explored transcriptomic differences between microglia morphologies, but these studies focused their assessments on the ’resting’ (ramified) vs. ’activated’ (ameboid) distinction to define morphology. It has yet to be determined whether transcriptomic profiles can be specified for the diverse morphologies such as rod-like and hypertrophic forms that lie outside of this dichotomy(26–28,62,63) and how these profiles relate to microglial function.

To begin to examine the relationship between different microglial morphologies and their potential spatial transcriptomic signatures, we developed a novel approach to combine different dataset modalities. As a proof of principle, we investigated the relationship between microglial morphology and gene expression in the human brain to define classes of microglia morphologies within a published Visium-SPG dataset(8). By leveraging a high-throughput morphology analysis toolset that we recently developed(29) and custom ImageJ and R scripts, we integrated the morphological information with the spatial coordinates of the barcoded spots in the Visium-SPG dataset. The coupling of these data types enabled us to link the spot-level data with the morphological classes of the microglia within the spots, while considering cell type composition and other spatially relevant variables. While the goal of this work was to develop a methodological approach to integrate morphology and spatial transcriptomics data, we were also able to identify several microglia morphology-relevant transcripts in our DE analysis that were consistent with previous work(6,25,44,57,58).

Until recently, it was not possible to perform untargeted whole-transcriptome profiling alongside morphological analysis from the same cells, limiting our ability to directly link cellular morphology to transcriptional state. Given that the identification of distinct, histologically defined microglial morphologies using Visium-SPG data has not yet been demonstrated, we first aimed to assess whether it would be possible to uncover the same morphology clusters commonly observed in other microglial studies in our analysis. We were able to identify four morphology classes - ameboid, hypertrophic, elongated, and ramified - previously characterized in a published mouse dataset(29). (Fig. 1G) Additionally, we were able to distinguish between different morphologies despite the Visium-SPG dataset being single plane 2D images, which was in line with prior work showing that relative group differences between different morphological classes are maintained across imaging parameters.(29) The morphological classes that we identified in our study exist along continuous scales of ramification (branching complexity), territory span, circularity (cell shape), and branch length(29,30). (Fig. 1A) Thus, individual features in isolation provide incomplete representations of distinct morphologies. For example, hypertrophic microglia differ from homeostatic, ramified microglia not only by their extent of ramification, but also by their larger soma size, increased thickness of branches, and reduced territory span, which together give them their characteristic ‘bushy’ appearance. (Fig. 1G,I) Although several studies have examined individual morphological features in relation to the expression of microglial cluster markers within the same cells(25,33), ours is the first to systematically link transcriptomic profiles to distinct morphological classes defined by multiple features together in a spatially-resolved, data-driven manner in the healthy human brain. Relating microglial gene expression and morphology will be invaluable to gain a more holistic view of microglia nomenclature, especially in disease contexts such as Alzheimer’s where morphology-relevant genes may change expression in relation to pathology.

Combining the immunofluorescent data with the spot coordinates also allowed us to explore microglia in comparison to other cell type signals in the dataset more thoroughly. This was important as microglia are known to have lower total mRNA content compared to other brain cell types.(64,65) While microglial marker genes were not correlated to TMEM119 fluorescent signal at the spot-level (Fig. S2A) nor amongst the highest expressed genes compared to other brain cell type markers (Fig. S2B-C), microglial marker genes were identified as medium-expressing across the final spots included in downstream DE analysis (Fig. S2B). This gave us confidence that although microglial genes were not the most highly expressed, true microglial signals could be identified when isolating and parsing spots specifically based on microglia morphology class.

Integration of the morphology and spot-level data allowed us to calculate novel spot-level measures that we considered along with cell type and white matter versus grey matter regional composition (Fig. 3B) in downstream DE modeling on pseudobulked samples created for each morphology class. 11 of the 34 total DEGs across comparisons overlapped with genes that are more highly expressed in human white matter microglia compared to grey matter microglia in both healthy and diseased conditions. While we observed previously identified microglia morphology-relevant genes(6) within our total DEGs, they overlapped highly with WM-related microglia genes. (Fig. 4C) Microglia share a close relationship with oligodendrocytes in both the developing(66,67) and adult,(58,68,69) brain, and the white matter has been shown to have a higher density of microglia compared to the gray matter in adult human brains(66,68,69). The incorporation of the regional composition effects in our statistical approach allowed us to explore potential state differences in microglia between white and grey matter. This differentiation may be particularly important for future applications examining microglia in the context of white matter diseases like MS which have a strong spatial localization of inflammation to the white matter lesions.

Across all DE contrasts (1 vs. all and pairwise), we observed the most DEGs in the ameboid form in comparison to all other forms. (Fig. 4A; Supplementary Table 2) We also identified two genes in this DE comparison, *SPP1* and *CD74*, which, while not meeting our criteria for statistical significance, have previously been extensively characterized in the literature as important markers of disease-associated microglial (DAM) states and related to microglia morphology differences in DAM.(25,25,57–59) DAM are distinct subsets of activated microglia linked to neurodegenerative diseases, characterized by a shift in gene expression related to inflammation, phagocytosis, and tissue repair.(25,70–72) *CD74* and *SPP1* were previously identified as cluster markers for human microglial subgroups that were specific to the immune activation axis and terminal DAM-like trajectory, respectively, and were uniquely more divergent from the homeostatic phenotype towards a DAM-like state compared to other clusters in a recent study from Tuddenham et al.(25). Ameboid microglia are commonly observed in postmortem brain tissue from patients with neurodegenerative disorders including Alzheimer’s and Huntington’s disease and display increased expression of various DAM-associated microglial genes including *CD74* and *SPP1*.(26,58,73–75) *CD74* has been found to accumulate in ‘reactive’ microglia, which are rounded in shape, in a rodent model of ischemia(57) and ameboid *Spp1*-expressing microglia have been observed at the periphery of tumors in glioma-bearing mice(58), together suggesting that these genes may play critical roles in defining microglia morphology in disease associated states.

While we highlight the novelty of our data analysis approach, we acknowledge several limitations. Expression profiles of individual Visium spots in the human DLPFC are from an average of 3 cells per spot(8,76). Thus, while we attempted to account for regional makeup and cell type composition, we were unable to directly link morphological data to transcriptomics data at the level of single microglia cells. However, no current methods exist that enable the direct coupling of IF with untargeted transcriptomics data at the single-cell level with detailed morphology. While spatial transcriptomics technologies like 10X Genomics’ Xenium *in situ* or VisiumHD platforms provide improved cellular resolution, they still present challenges in relating the exact cell types to corresponding signals and gene panels need to be designed *a priori* for Xenium. Multiplexed immunofluorescence panels also present similar limitations, as they require prior knowledge of the markers to be studied. In contrast, the data type analyzed in our work, Visium-SPG, while limited in cellular resolution, captures the whole transcriptome of each barcoded spot and thus does not require prior knowledge of morphology-relevant genes. While VisiumHD, the newest iteration of 10X Genomics spatial gene expression technologies, allows for subcellular sampling (2um bins) which positions it as a good candidate for reconstruction of whole cell morphology *in situ*, the assay would require further optimization of immunofluorescence staining from the same tissue to enable segmentation of microglia morphology. Finally, our study is limited by its small sample size of 3 human subjects and 1104 final Visium spots after application of our stringent filtering criteria (Fig. 2A). While we were able to identify morphology-relevant microglial genes in our analysis by filtering, there were likely additional non-microglial cell type signals and regionally-driven genes that were present in the pool of final DEGs. Larger scale datasets in the future may yield more morphology-specific signals, especially in disease contexts where significant shifts in microglia morphology are anticipated in relation to pathology. Future studies should also allow for well-powered analysis by sex, an important factor for understanding microglia function(77).

Despite these challenges, our method remains valuable and provides broader utility, and the analysis approach presented in this work can be applied to other datasets and experimental contexts. Technological advancements in spatial biology will continue to yield more datasets of multiple dimensions including morphological and transcriptomic data, emphasizing the need to develop new approaches to integrate these different layers of information. In future studies, we envision the application of our integrative approach and method in new ways that offer a deeper understanding of microglial roles across different biological contexts. This will include characterizing spatial distributions of morphological classes in relation to specific cell types and in disease states like Alzheimer’s and MS, which have important spatial qualities to their pathology.

## Data availability

All analysis code and processed data can be found on the Ciernia Lab Github at https://github.com/ciernialab/Kim2026_VisiumMicrogliaMorphology and on the Open Science Framework (OSF) website at https://osf.io/uayfx.

## Acknowledgements

We thank the Ciernia Lab and Pavlidis Lab members for their thoughtful feedback and suggestions during lab meetings throughout the progression of this project. We are grateful for the resources provided by the Neuroimaging & Neurocomputation Centre and the Dynamic Brain Circuits in Health & Disease Research Cluster at the University of British Columbia.

## Figures

**Figure S1.**
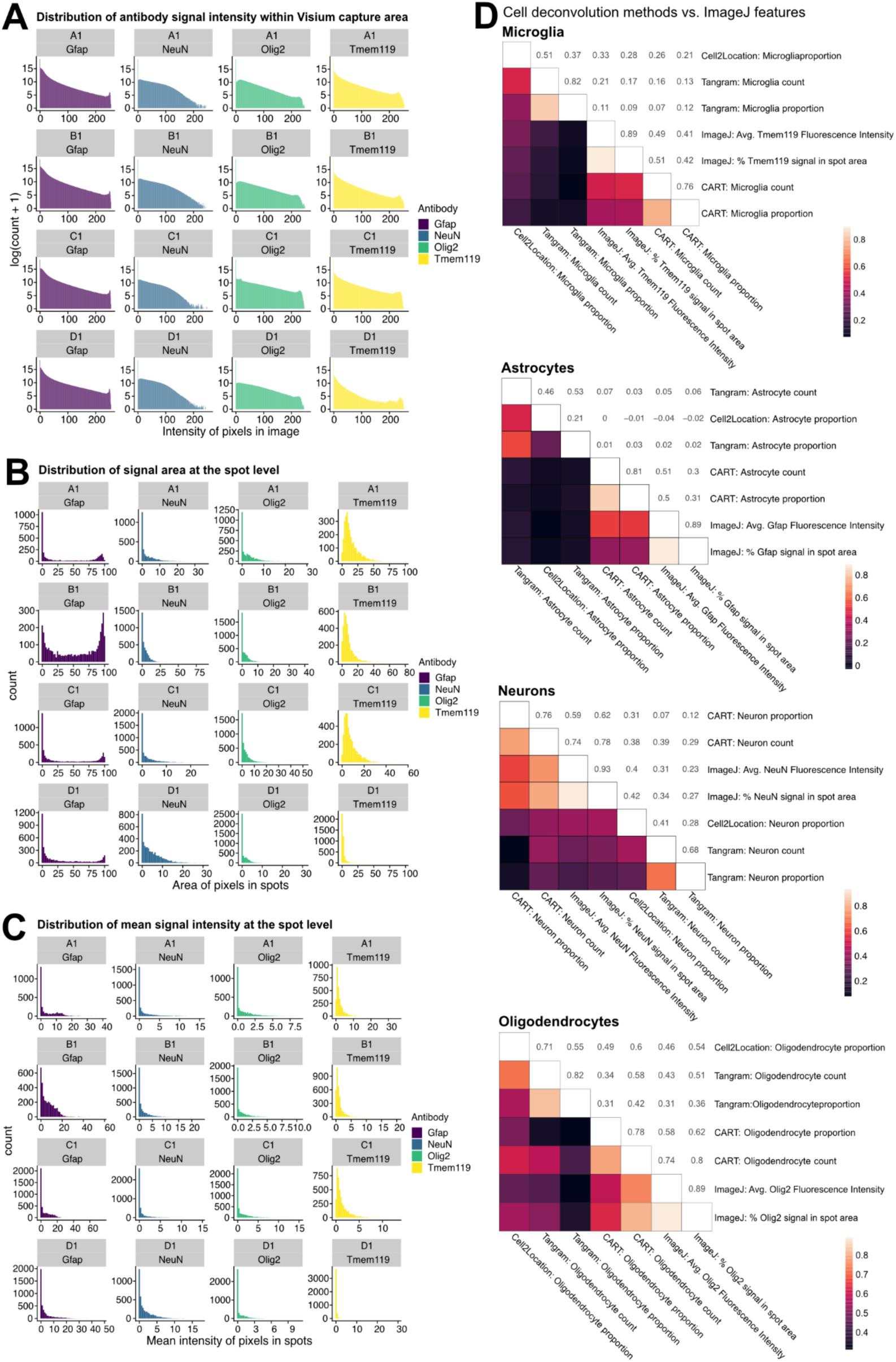
ImageJ exploration of GFAP, NEUN, OLIG2, and TMEM119 channel immunofluorescent signal in Visium-SPG dataset. (**A**) Distribution of mean channel signal intensity of each marker within larger Visium capture area for each Visium-SPG image in dataset. (**B**) Distribution of channel signal area within each of the Visium spots for each image in the dataset. (**C**) Distribution of mean channel signal intensity within each of the Visium spots for each image in the dataset. (**D**) Pearson’s correlation of ImageJ-derived features (cell type-specific channel mean signal intensity, percentage of spot area taken up by signal) and cell deconvolution counts for each cell type. *CART* estimates are based on the immunofluorescent imaging data, and *Tangram* and *Cell2location* infer cell types based on gene expression.

**Figure S2.**
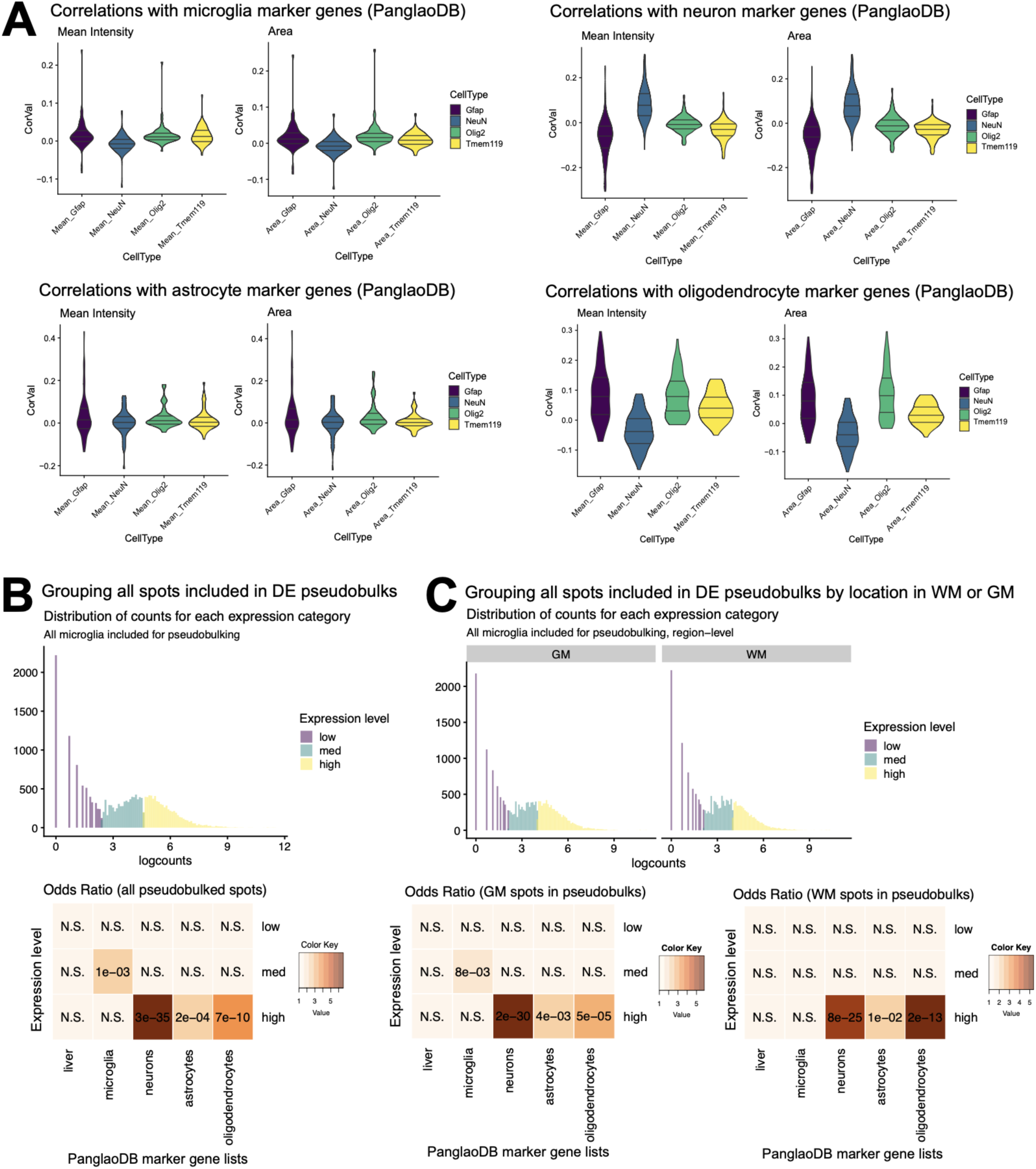
Exploring cell type-specific gene expression within Visium-SPG dataset. (**A**) Correlations of expression of cell type-specific marker genes from PanglaoDB database(7) with ImageJ-derived features (channel mean signal intensity within Visium spot, percentage of spot area taken up by channel signal) for each cell type. (**B**) Distribution of counts for low, medium, and high-expressing genes and their overlap with PanglaoDB marker gene lists for different cell types after grouping all spots included in DE pseudobulks. Odds ratio values represented by color and BH-adjusted p-values overlaid. (**C**) Distribution of counts for low, medium, and high-expressing genes and their overlap with PanglaoDB marker gene lists for different cell types after grouping all spots included in DE pseudobulks within white matter and grey matter separately. Odds ratio values represented by color and BH-adjusted p-values overlaid.

**Figure S3.**
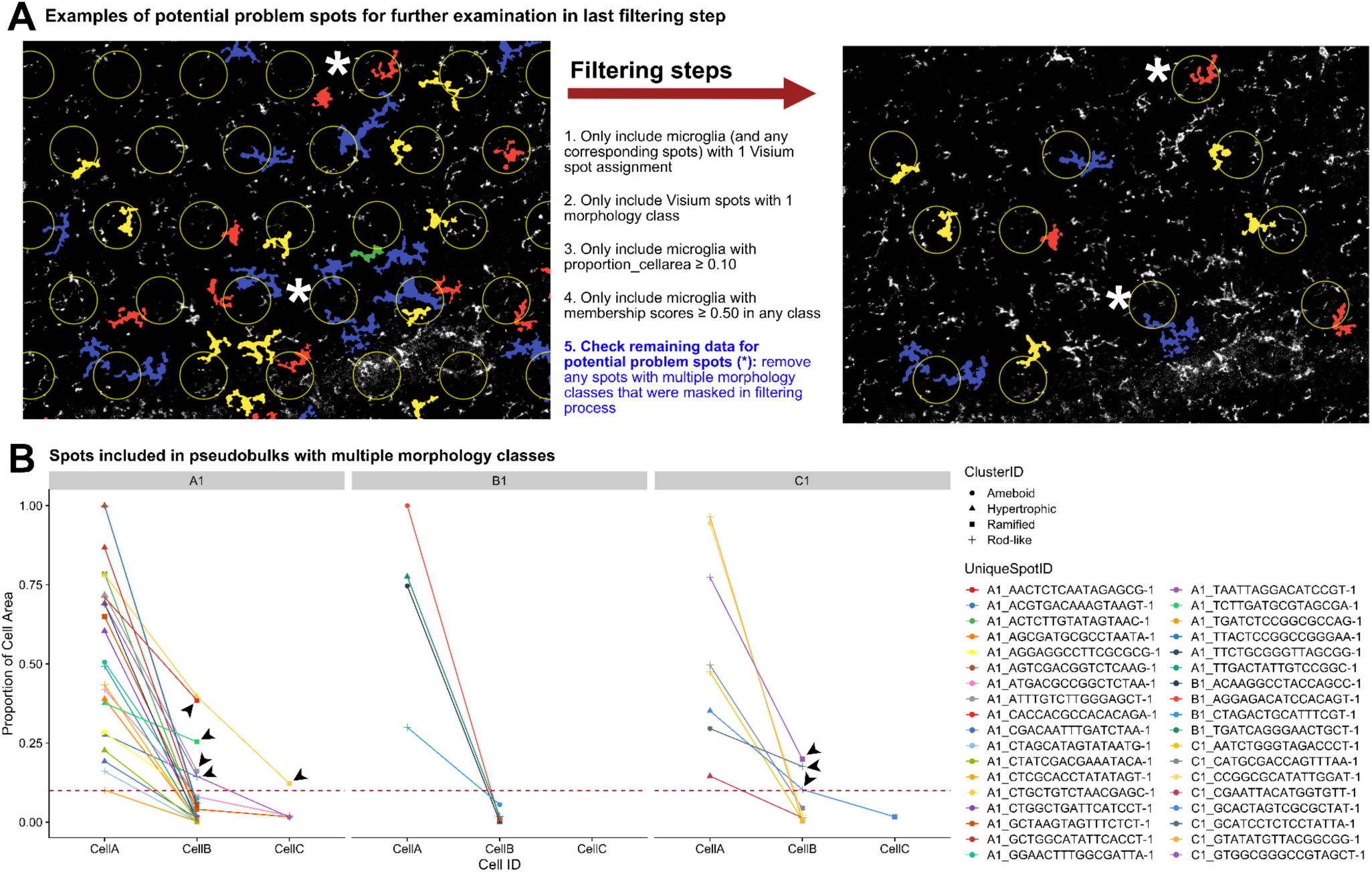
Evaluation of potential problem spots after application of spot filtering criteria. (**A**) Example visualization of 2 potential problem spots (denoted by *) that in reality had >1 morphology classes present (top *) or >1 microglia (bottom *), properties that were masked in the filtering of the integrated dataframe shown in Fig. 2A. (**B**) Evaluation of potential problem spots that remained after steps 1-4 of filtering criteria. To stay faithful to the original filtering criteria, spots were removed only if they contained multiple microglia with proportion_cellarea ≥ 0.10 that were of different morphology classes. Dotted line denotes proportion_cellarea = 0.10 and arrowheads point to Visium spot IDs that were removed.

**Figure S4.**
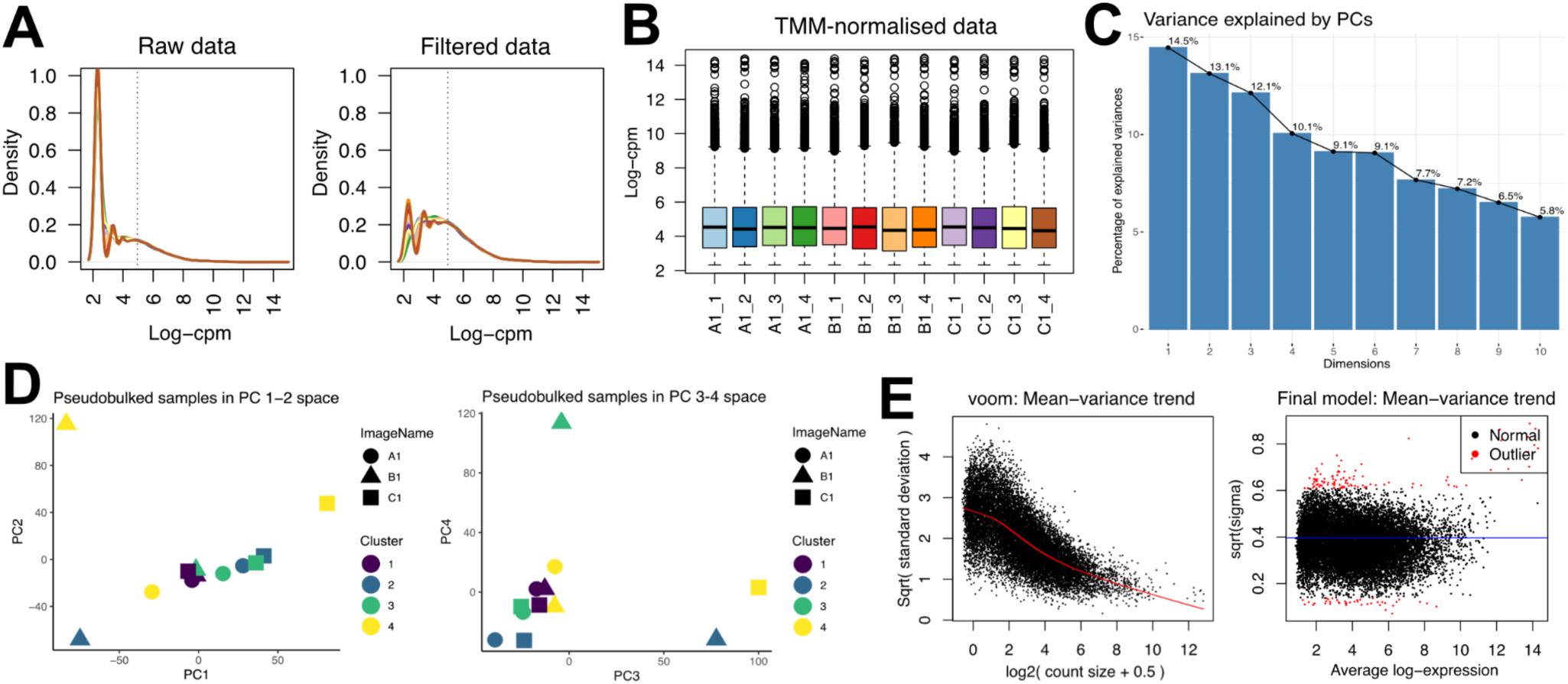
Differential gene expression analysis using limma-voom. (**A**) Distribution of log2CPM for each pseudobulked sample before and after filtering for zero counts. (**B**) Log2CPM values for each pseudobulked sample after TMM normalization. (**C**) Variance explained by first 10 PCs after dimensionality reduction by PCA. (**D**) Pseudobulked samples in PCs 1-3 space, colored by morphology class and denoted in shape by image name. (**E**) Mean-variance trend of counts and final statistical model after limma-voom.

**Figure S5.**
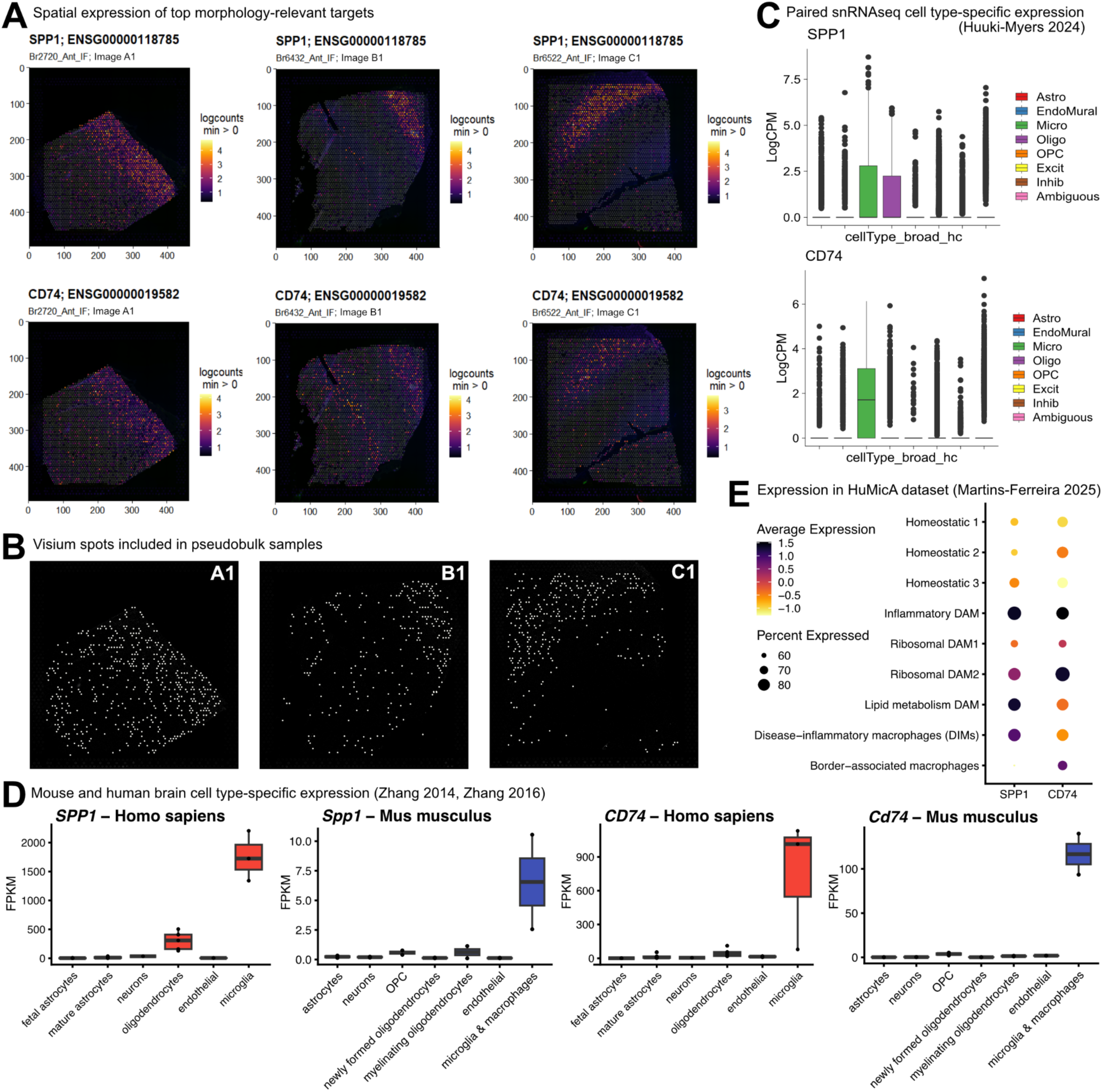
Example of microglial genes that are under threshold (SPP1 and CD74). (**A**) Spatial expression of *SPP1* and *CD74* plotted in DLPFC Visium-SPG images using SpatialLIBD R package. (**B**) Spatial distribution of Visium spots included for generation of pseudobulked samples, filled in white. (**C**) *SPP1* and *CD74* cell type-specific expression in paired snRNAseq data from Huuki Myers et al., 2024(8). (**D**) *SPP1* and *CD74* cell type-specific expression in human and mouse brain single cell atlases from Zhang et al., 2014(9) and 2016(10) (**E**) *SPP1* and *CD74* enrichment (percentage) and relative average expression (scaled across clusters) in Human Microglia Atlas microglia clusters from Martins-Ferreira 2025(11).

**Figure S6.**
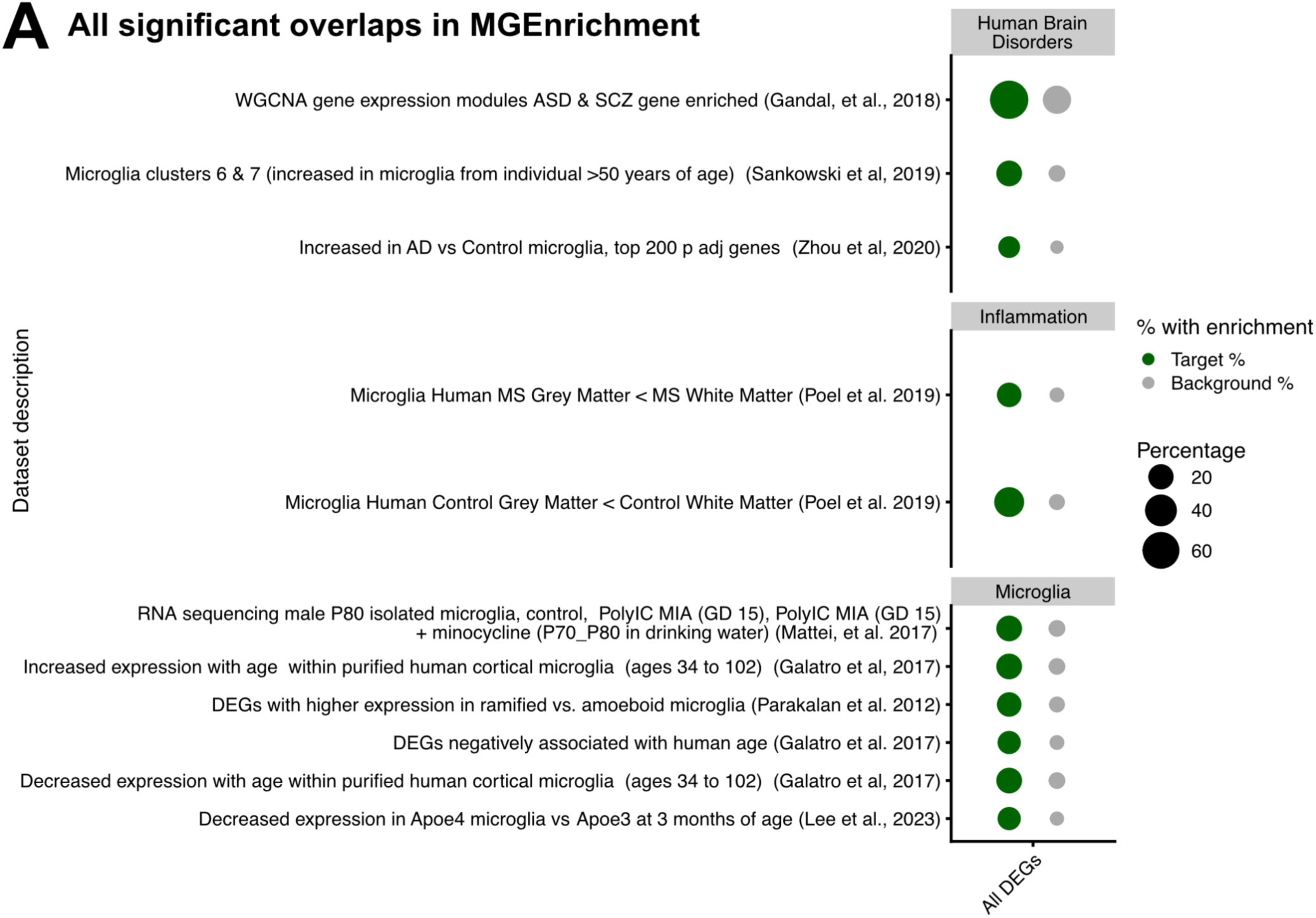
Significant DEG overlaps with published microglia gene lists. (**A**) Gene list overlap enrichment against publicly available microglia lists in MGEnrichment(4) and all 34 significant DEGs across all contrasts. Green circles denote a significant enrichment in the percent of DEGs for a given gene list compared to that of the background of all genes detected in the RNA-seq experiment (grey circles). Only significant enrichments are shown (BH-corrected p<0.05).

